# ezQTL: A Web Platform for Interactive Visualization and Colocalization of Quantitative Trait Loci and GWAS

**DOI:** 10.1101/2022.03.08.483491

**Authors:** Tongwu Zhang, Alyssa Klein, Jian Sang, Jiyeon Choi, Kevin M Brown

## Abstract

Genome-wide association studies (GWASs) have identified thousands of genomic loci associated with complex diseases and traits, including cancer. The vast majority of common trait-associated variants identified via GWAS fall in non-coding regions of the genome, posing a challenge in elucidating the causal variants, genes, and mechanisms involved. Expression quantitative trait locus (eQTL) and other molecular QTL studies have been valuable resources in identifying candidate causal genes from GWAS loci through statistical colocalization methods. While QTL colocalization is becoming a standard analysis in post-GWAS investigation, an easy web tool for users to perform formal colocalization analyses with either user-provided or public GWAS and eQTL datasets has been lacking. Here, we present ezQTL, a web-based bioinformatic application to interactively visualize and analyze genetic association data such as GWAS and molecular QTLs under different linkage disequilibrium (LD) patterns (1000 Genomes Project, UK Biobank, or user-provided data). This application allows users to perform data quality control for variants matched between different datasets, LD visualization, and two-trait colocalization analyses using two state-of-the-art methodologies (eCAVIAR and HyPrColoc), including batch processing. ezQTL is a free and publicly available cross-platform web tool, which can be accessed online at https://analysistools.cancer.gov/ezqtl.

## Introduction

Genome-wide association studies (GWASs) have proven powerful in identifying common genetic variants and loci associated with many complex diseases and traits, including cancer susceptibility [1]. Lead single nucleotide polymorphisms (SNPs) identified via GWAS are not necessarily causal variants themselves; rather these variants often tag a larger set of many variants in high linkage disequilibrium (LD), one or more of which may be causal variants underlying the trait [2]. Most of these trait-associated variants are non-coding, and for a large proportion of loci, causal variants are thought to be *cis*-regulatory. Expression quantitative trait locus (eQTL) studies have shown that common *cis*-regulatory variants often regulate many genes other than the nearest one [3], meaning that many distant genes must be considered plausible candidate genes. Sifting through and functionally characterizing a large number of statistically-correlated candidate variants, as well as in many cases identifying the likely causal gene(s) affecting disease risk at GWAS loci, remain significant challenges in the field. These challenges hinder the translation of GWAS findings to a better understanding of the biological processes underlying complex traits including cancer susceptibility.

Most common causal risk variants are hypothesized to function by affecting gene expression, or alternatively, via potentially affecting patterns of splicing, DNA methylation, and/or expression levels of microRNAs or other non-coding RNAs. Genetic association tests of these various molecular quantitative trait loci (QTLs) are commonly used for prioritization of common variants within GWAS loci for functional study, as well as identification of the gene(s) likely to underlie risk at these loci [4–6]. Numerous eQTL datasets derived from normal human tissues have been made publicly available, such as the Genotype-Tissue Expression (GTEx) Project [7], which has become a key resource for investigating tissue-specific QTLs underlying complex traits. Colocalization analysis has been proposed as an important step to test whether complex trait associations (for example, cancer risk) and molecular QTLs (such as those for gene expression, eQTLs) share common causal genetic variants for both signals, which will aid biological insight following GWAS studies. Statistical colocalization methods have been developed to assess the overlap of genetic association signals across multiple related traits (such as molecular QTLs and GWAS) [8]. Rather than focusing on lead variants, a colocalization analysis compares the distribution of summary statistics from two association signals and accounts for LD, which reduces the false positive discoveries by using multiple variants [9–11]. However, current web-based applications, such as LocusZoom [12], do not provide visualization of GWAS-QTL colocalization, whereas LocusCompare [13] provides only limited colocalization statistics. Although there is an abundance of implementation of colocalization algorithms in R and tools for visualization, running these various programs may require additional QC of user-supplied data and different data formatting, which can be challenging for a wet-lab researcher with little background in bioinformatics. There is also a critical need for a colocalization platform with large available QTL datasets including different types and tissues, and LD datasets including different populations.

To address this problem, we developed ezQTL to interactively visualize the genetic associations and perform colocalizations between two traits. The web-based modules (Locus QC, Locus LD, Locus Alignment, Locus Colocalization, Locus Table, Locus Quantification, and Locus Download) utilize GWAS summary statistics, QTL association data, and LD matrix data to perform comprehensive colocalization analysis, culminating in a quality control (QC) report and interactive visualization. ezQTL bridges current gaps of existing colocalization analysis tools by integrating user-supplied as well as publicly available GWAS and QTL resources, LD reference datasets, different colocalization algorithms, and interactive visualization, through a user-friendly web interface.

### Implementation

The ezQTL web application includes seven modules (**Figure 1**): Locus QC, Locus LD, Locus Alignment, Locus Colocalization, Locus Table, Locus Quantification, and Locus Download. All seven ezQTL modules are written in R and run on a virtual machine using the UNIX operating system. The webtool is primarily written in Javascript and R. All web content is programmed in HTML5 for cross platform versatility. ezQTL is built using the following packages for colocalization analysis: eCAVIAR (https://github.com/fhormoz/caviar) and HyPrColoc (https://github.com/jrs95/hyprcoloc). It also uses the IntAssoPlot for LD visualization (https://github.com/whweve/IntAssoPlot). The Plotly R graphing library is used to generate interactive plots. All datasets, QC files, and results, including plots and tables in each analysis module, can be downloaded using the Locus Download module. For genomic data privacy, the input data in ezQTL is encrypted and secured during the data transition, calculation, and temporary storage. All input data and results for each query or submission will be deleted automatically after seven days.

**Figure 1.**
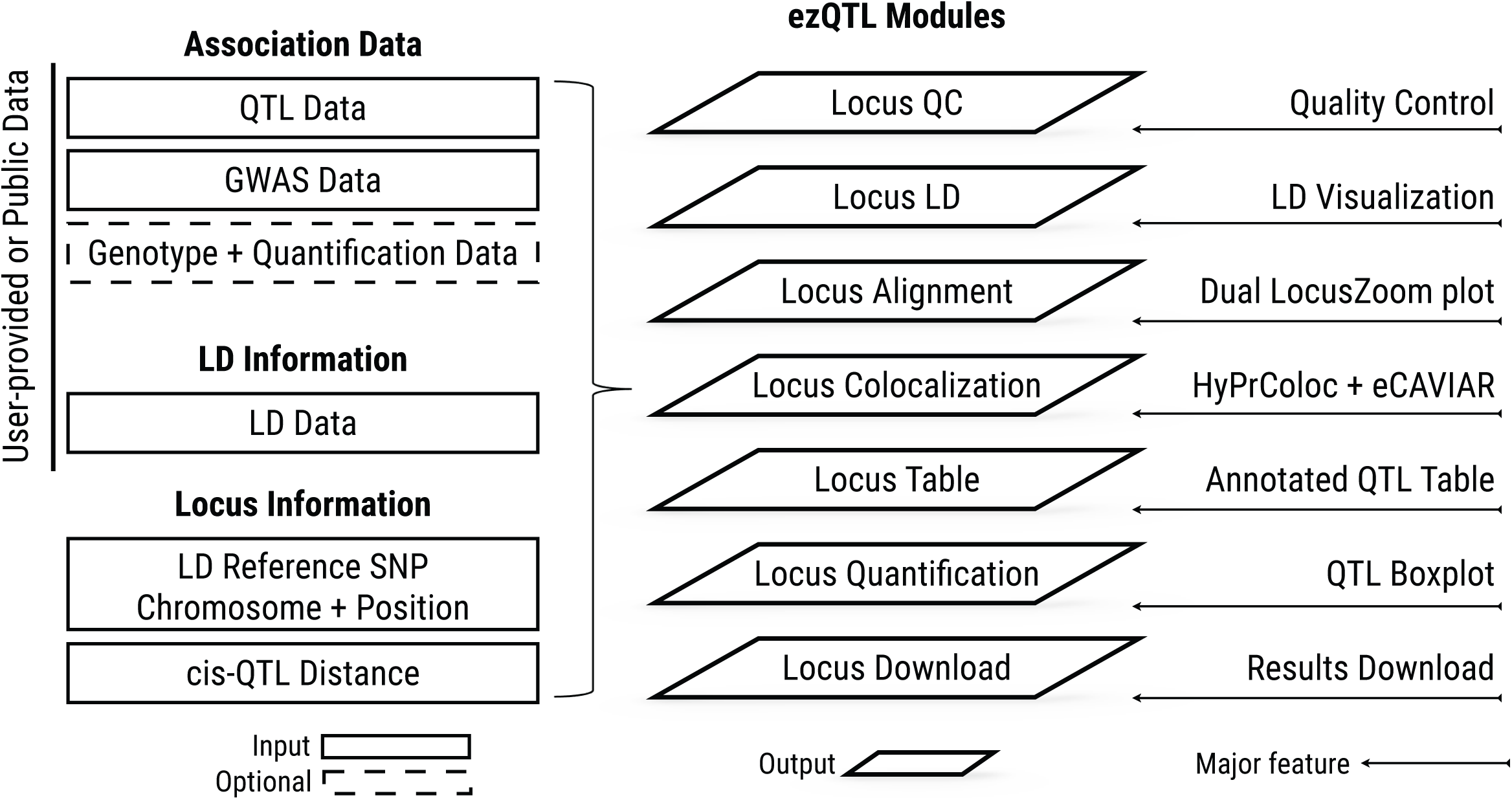
The architecture of the ezQTL analysis platform including the input data types, analysis modules, and major features are shown.

### User- and publicly-supplied input data

Currently, for colocalization, ezQTL requires QTL, GWAS, and LD matrix data that can be supplied by the user. However, ezQTL also hosts a large collection of publicly available QTL, GWAS, and LD datasets, currently 594 datasets collectively listed on the searchable Public Data Source tab. We provide public datasets that are pre-processed into the standard ezQTL input format and example scripts to assist users in preparing user-supplied datasets into the correct input format. GWAS summary statistics included in ezQTL can be downloaded at https://github.com/mikegloudemans/gwas-download and https://www.ebi.ac.uk/gwas/home. The GTEx datasets for both eQTL and sQTL can be accessed via https://gtexportal.org/home/datasets. The link used to download other GWAS or QTL datasets from individual studies can be accessed through the ezQTL website. 1000 Genomes data used to generate the LD matrix are available at http://ftp.1000genomes.ebi.ac.uk/vol1/ftp/release/20130502/. The LD matrix from the UK Biobank dataset can be accessed at https://storage.googleapis.com/broad-alkesgroup-public/UKBB_LD.

User-supplied QTL summary statistics can be provided for any given locus to be tested (e.g. by default variants +/− 75 kb of the lead SNP can be used) in the same format as the output from commonly used QTL data analysis tools (e.g. FastQTL [14] and Matrix eQTL [15]); the fields required by ezQTL are matched by the column header names of the input file and a warning is provided when no match is found. ezQTL also provides commonly used QTL datasets, including both sQTLs and eQTLs from GTEx in two genome builds (GRCh37 and GRCh38) [16] and QTLs from additional tissue types [4,17–23].

For GWAS summary data, ezQTL similarly accepts user-supplied summary data in a general format for each locus (e.g., variants +/− 75bp of the lead SNP), matched by column headers for the required fields. We also provide an example script to transfer GWAS summary statistics from NHGRI GWAS catalog into ezQTL GWAS data format through ezQTL Github page. Currently, ezQTL has pre-loaded GWAS summary statistics previously collected by LocusCompare [13] and most of the publicly available cancer-related GWAS summary data downloaded from the NHGRI GWAS Catalog [24] or the study websites. At this time, there are more than 420 phenotypes included in the set of pre-supplied GWAS summary data.

For LD reference data, ezQTL will instantly calculate the LD matrix for any locus from any combination of 1000 Genomes (Phase 3) populations [25] for both GRCh37 and GRCh38 genome builds, or alternatively provide a pre-calculated LD matrix from the UK Biobank European populations [26] (GRCh37 only). Users may also supply their own LD matrix data generated by emeraLD [27].

In addition to colocalization analysis using these three data types, users can provide raw QTL data (e.g., matched gene expression or other QTL trait levels and individual genotypes) for extended data QTL visualization functions. We anticipate future implementations of this visualization feature extended to pre-loaded public eQTL data such as those from GTEx.

### Features of ezQTL modules

ezQTL takes as minimal input both GWAS and QTL data for colocalization functions and any one of GWAS, QTL or LD for visualization functions. The detailed relationship between module functions and input datasets are included in the documentation of ezQTL. The Locus QC module performs data quality control for user-supplied GWAS, QTL, and or LD data (such as for data formatting, allele-matching, variant filtering and suggesting a reference variant) and generates a variant level summary report for each dataset and reports overlapping SNPs. This module also provides visualizations for each input dataset and highlights potential issues (such as insufficient overlapped variants) for subsequent colocalization analysis. The post-QC data generated by Locus QC will be used for all other modules.

Locus LD and Locus Alignment modules both visualize QTL and GWAS data together with LD information for the locus to provide a quick visual overview of the colocalization pattern. Locus LD generates Manhattan plots for either QTL or GWAS *P*-values along with local LD patterns using the IntAssoPlot R package [28]. The Locus Alignment module simultaneously and interactively visualizes association *P*-values and linkage disequilibrium patterns (recombination rate) for both GWAS and QTL datasets using two LocusZoom plots for a given gene or probe in a locus of interest. An example data is provided for the melanoma GWAS locus at Chr21q22.3 and melanocyte-specific eQTL data for *MX2* [4,29] (**Figure 2**). In addition, clicking on any given variant on the LocusZoom plot can directly connect to other functions, such as setting that variant as the LD reference, creating a QTL boxplot (if users provide individual level QTL data), or linking to multiple external databases (LDpop [30], GWAS catalog [24], gnomAD [31]). A *P*-value correlation plot between these two LocusZoom plots, similar to that provided by LocusCompare, is generated to visualize colocalization.

**Figure 2.**
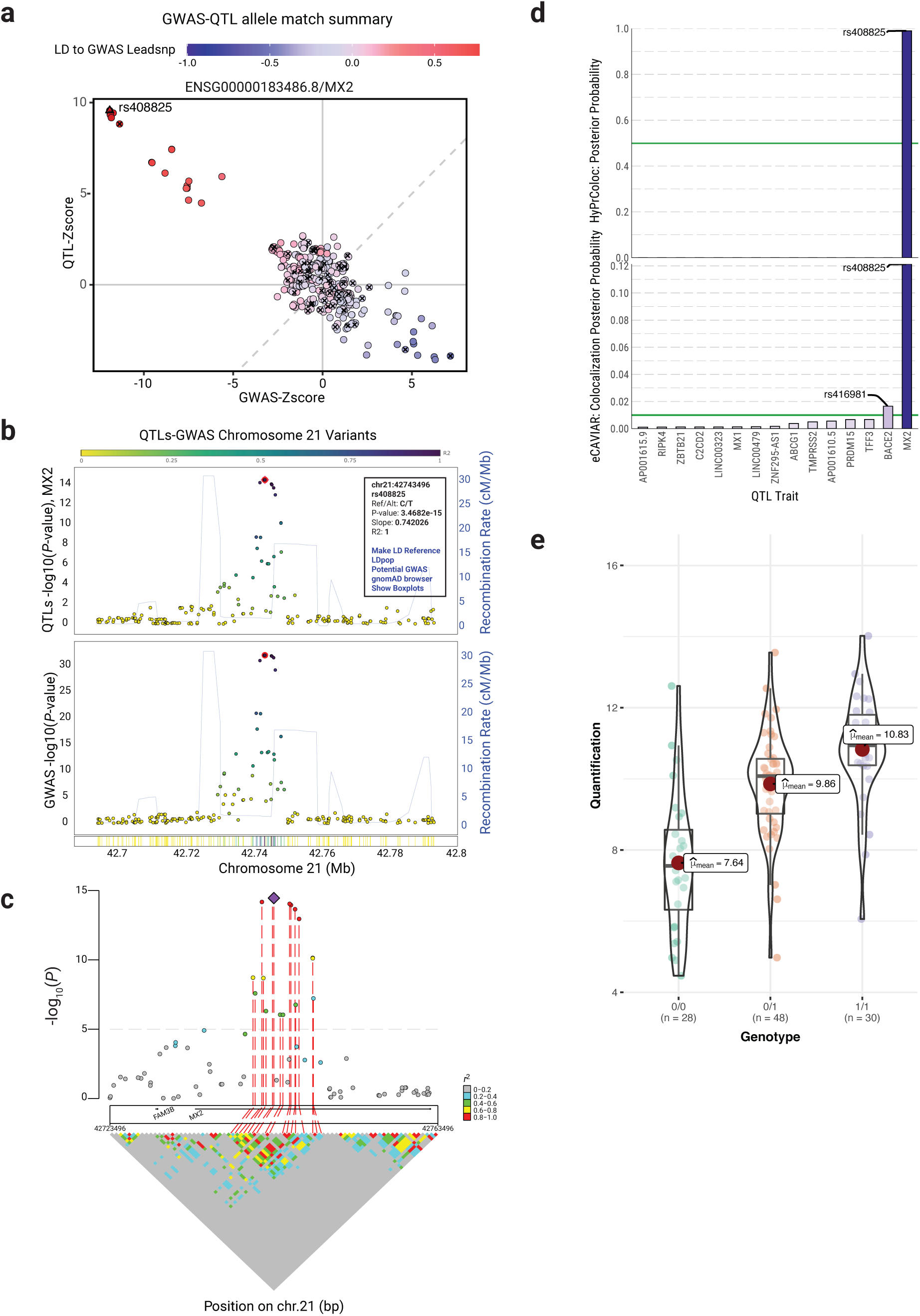
Example of ezQTL colocalization analysis between GWAS and eQTL in a melanoma GWAS locus. **A**. Correlation plot of Z-scores between QTL and GWAS from the Locus QC module for identification of potentially mismatched alleles. Variants marked by “x” are C/G or A/T SNPs. **B**. A dual LocusZoom plot with eQTL association on top and GWAS association on the bottom from Locus Alignment. The pink diamond in the LocusZoom plot indicates the most significant SNP among the eQTL association, and the red circle highlights the current LD reference SNP. The pop-up window after clicking each SNP shows association information, additional actions, and links. **C**. LD matrix visualization from Locus LD. The pink diamond and red dashed line indicate the reference SNP. **D**. Colocalization analysis by both eCAVIAR and HyPrColoc from Locus Colocalization. **E**. QTL boxplot between genotype of reference SNP and gene quantification from Locus Quantification. The statistical method and significance are labeled on the bottom and top of the boxplot, respectively. QTL, quantitative trait locus; GWAS, genome-wide association study; QC, quality control; SNP, single nucleotide polymorphism; eQTL, expression quantitative trait locus, LD.

The Locus Colocalization function performs colocalization analysis between GWAS and molecular QTL (or any two traits) within a locus using two different algorithms, eCAVIAR [9] and HyPrColoc [8] (**Figure 2**). By default, data used for colocalization is selected from a genomic window centered on the variant with the smallest GWAS *P*-value within the region uploaded. Re-centering on a different LD reference SNP can be specified either at the time of uploading the data, or alternatively by entering or selecting a specific variant on the LocusAlignment page following data upload. HyPrColoc [8] is a computationally efficient method based on the Bayesian algorithm to estimate the posterior probabilities that a variant is causal in both GWAS and QTL studies. This approach extends the underlying methodology of COLOC [32] and MOLOC [33]. As implemented in ezQTL, HyPrColoc uses all genetic variants within a user-specified window (specified on the data input panel, the default is +/− 75kb centered over the lead GWAS or a user-specified reference variant). For eCAVIAR, the colocalization test is performed by multiplying posterior causal probabilities from the QTL and GWAS studies, to generate a colocalization posterior probability (CLPP), as described by Hormozdiari *et al*. [9]. eCAVIAR colocalization as implemented in ezQTL is performed both using all genetic variants within the user-specified window used by HyPrColoc, as well as simultaneously for the set of 100 variants surrounding the lead GWAS variant (50 variants on each side as well as the lead variant itself). Both eCAVIAR and HyPrColoc results are presented and compared as exportable barplots and tables including all the molecular traits (such as eQTL genes) tested in each locus. We recognize that the colocalization results for two traits in the same locus might be different based on different colocalization approaches. HyPrColoc is a deterministic Bayesian colocalization algorithm that makes the assumption of a single causal variant in a given trait, similar to other existing methods, such as COLOC [32] and MOLOC [33]. Although HyPrColoc can be used for multiple-trait colocalization, the HyPrColoc analysis implemented in ezQTL is set for two-trait analysis at a time and does not use linkage disequilibrium (LD) information (except for two traits with overlapped samples) because of the single causal variant assumption. eCAVIAR is a probabilistic model that could consider multiple causal variants in a locus, and the eCAVIAR analysis provided by ezQTL is allowing up to two casuals using the LD information that users choose for the analysis. Some loci may harbor more than one causal variant for any given trait, and the colocalization results under the single causal variant assumption would not hold when secondary causal variants explain as much trait variation as the identified common causal variant [8]. On the other hand, eCAVIAR was designed to account for allelic heterogeneity and LD, under the assumption that the number of individuals and the LD is the same between GWAS and QTL populations [9]. However, misspecification of the LD structure when aiming to fine-map more than a single causal variant can lead to major biases in the results. If an accurate LD reference panel of a large sample size that matches both GWAS and QTL populations is not available, a single causal variant model without using LD could be more reliable [8]. In addition, the number of variants selected for colocalization analysis are different between eCAVIAR and HyPrColoc, as described in the Locus Colocalization module. Thus, we recommend users to consider these differences in the statistical models and assumptions as described in the original publications of eCAVIAR [9] and HyPrColoc [8] when reporting and interpreting the results.

Locus Quantification provides visualizations (violin, distribution and pairwise correlation plots) of optional quantification data used in the QTL testing of the locus of interest. The module requires user-provided individual-level genotypes and molecular trait data that were used for calculation of QTLs (e.g., gene expression or DNA methylation); in the current version, QTLs provided by ezQTL are pre-computed summary statistics and do not use individual level genotype or trait data. These data allow generation of violin plots for each trait, for example, in the Locus Quantification module, as well as boxplots for QTLs that can be accessed through the Locus Alignment module. Finally, Locus Table provides a sortable table of user-provided QTLs for any given trait in a region. QTL results are annotated with linkage disequilibrium to any given index SNP and links to multiple external databases. All datasets, QC files, and results, including plots and tables in each analysis module, can be exported using the Locus Download module.

### Comparison with other tools

In comparison to similar tools (such as LocusZoom [12], LocusCompare [13], LocusFocus [34], GTEx Portal [35] and eQTLplot [36]), ezQTL contains several new features and functions for integrated analysis of GWAS and QTLs (**Table 1**). ezQTL provides interactive dual-LocusZoom plots, which allows users to explore and integrate the association data between GWAS and QTL with availability to recalculate LD information based on selected reference variant. ezQTL performs variant level QC before colocalization analysis for all user-supplied and pre-provided datasets, and also generates reports and plots for future investigation. For example, matching allelic directions between GWAS and QTL datasets is essential for colocalization analysis, and ezQTL will report any flipped or unmatched variants to users for recalibration. ezQTL hosts a large number of public datasets (including those from different genome builds, different tissue types and different populations) for each data type and allows users to perform colocalization analysis using mixed data sources. These public datasets in ezQTL also allow users to perform exploratory data analysis in any given locus. ezQTL also implements two state-of-the-art methodologies (eCAVIAR and HyPrColoc) for formal colocalization analysis, which represent two distinct statistical approaches with different assumptions of the number of causal variants. In addition, serial visualizations have been implemented in ezQTL, including multiple QC related plots, p-value correlation plots, LD matrix visualization, colocalization summary plots, and QTL association plots. For the same dataset, ezQTL is designed to allow users to perform analysis for multiple loci by inputting different locus information. In addition, ezQTL allows users to submit a job to the ezQTL server and an email with a link to access all of the results will sent to the user.

**Table 1.**
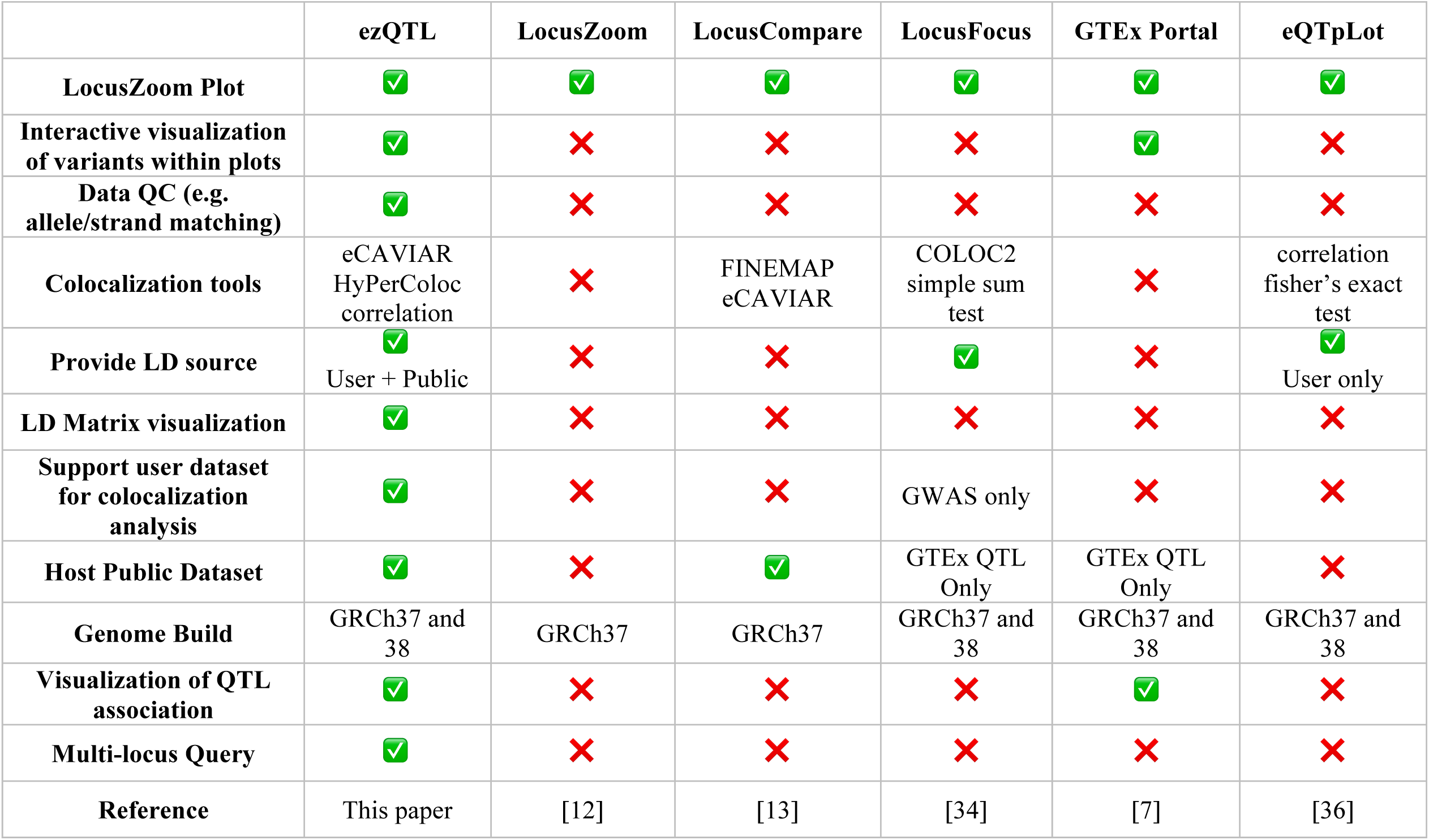

## Conclusions

Here, we present ezQTL as a web tool to interactively visualize and analyze genetic association data and perform colocalizations between two traits without much bioinformatics skill. ezQTL is competitive with two state-of-the-art colocalization approaches as well as large public datasets. ezQTL enhances existing colocalization analysis tools for GWAS study by integrating user-supplied as well as publicly available GWAS and QTL resources, LD reference datasets, different colocalization algorithms, and interactive visualization through a user-friendly web interface. In summary, ezQTL facilitates mapping disease susceptibility regions and assist researchers in characterizing and prioritizing functional genes and variants based on the genotype-phenotype associations.

## Code availability

ezQTL is a free and publicly available cross-platform web tool which can be accessed online at https://analysistools.cancer.gov/ezqtl. The source code for ezQTL can be found at CBIIT GitHub page: https://github.com/CBIIT/nci-webtools-dceg-ezQTL and at BioCode at NGDC: https://ngdc.cncb.ac.cn/biocode/tools/BT007295. It is licensed under the GNU General Public License version 3 (GPLv3).

## CrediT authorship contribution statement

**Tongwu Zhang**: Conceptualization, Software, Resources, Data Curation, Writing – Original Draft, Visualization, Funding acquisition. **Alyssa Klein**: Software, Validation, Data Curation. **Jian Sang**: Data Curation, Validation. **Jiyeon Choi**: Conceptualization, Validation, Resources, Data Curation, Writing – Original Draft, Supervision, Funding Acquisition. **Kevin Brown**: Conceptualization, Validation, Resources, Data Curation, Writing – Original Draft, Supervision, Funding Acquisition.

## Acknowledgments

The authors thank Kailing Chen, Kevin Jiang, Phyllip Cho, and Ben Chen from the National Cancer Institute Center for Biomedical Informatics and Information Technology (CBIIT) for technical assistance and web development, as well as the National Cancer Institute’s Laboratory of Translational Genomics and Division of Cancer Epidemiology and Genetics for valuable input and testing. Support comes from the National Cancer Institute’s Intramural Research Program (1ZIACP010201) and the Division of Cancer Epidemiology and Genetics Informatics Tool Challenge. The content of this publication does not necessarily reflect the views or policies of the US Department of Health and Human Services, nor does mention of trade names, commercial products, or organizations imply endorsement by the US government.

